# Diazepam alters the shape of alpha oscillations recorded from human cortex using EEG

**DOI:** 10.1101/2025.10.24.684311

**Authors:** George M Opie, Natalie Schaworonkow, Pedro Gordon, Dania Humaidan, Ulf Ziemann

## Abstract

While neural oscillations are conventionally assessed via their frequency, power and phase, developing literature suggests that their shape also provides neurophysiological and functional information. However, the extent to which the shape of oscillations recorded non-invasively in humans index specific brain processes remains unclear. This study implemented a pharmaco-EEG approach to begin addressing this limitation. In 21 healthy adults, resting-state EEG data was collected before and after placebo or diazepam, a positive allosteric modulator of type A γ-aminobutyric acid (GABA_A_) receptors. The shape of individual cycles in the alpha band was then derived using empirical mode decomposition, followed by extraction of principal components (PCs) describing specific facets of alpha shape. Results of this approach show that all shape features were unchanged following placebo. In contrast, diazepam was associated with complex changes in several shape features, including peak-trough shape and edge speed. While changes in shape were apparent in all cortical lobes, the strongest alterations were specific to sensorimotor and parietal cortices. Taken together, our results support the neurophysiological utility of waveform shape, particularly with respect to non-invasive human recordings. Furthermore, the regional specificity of effects highlights the need for more granular exploration of waveform diversity.

## Introduction

Neural oscillations represent a stereotypical feature of the brain’s electrical activity (Buzsáki and Draguhn 2004). These rhythmic fluctuations in voltage occur across a broad frequency range and are ubiquitous throughout the brain. Furthermore, they have been associated with critical brain functions such as inter-areal communication and signal integration (Fries 2005, Jensen and Colgin 2007, Fries 2015), and therefore underpin everyday processes like learning and memory (Buzsaki 2006, Canolty and Knight 2010, Buzsáki and Watson 2012). Alterations in oscillatory activity have consequently been associated with numerous pathologies and disorders, including schizophrenia (Uhlhaas and Singer 2010), ADHD (Calderone *et al*. 2014) and Parkinson’s disease (Oswal *et al*. 2013). Given the above, it is unsurprising that the examination of neural oscillations represents the focus of a huge literature. From a methodological perspective, this has involved an expansive suite of approaches. However, derivation of oscillatory power via Fourier-based techniques has been the mainstay; this approach has provided a powerful means to investigate complex oscillatory activity in many different contexts, and has driven countless neuroscientific discoveries.

While spectral decomposition via Fourier-based methods captures some important features of neural oscillatory activity, it disregards others. In particular, the underlying assumption that a sinusoidal waveform best replicates endogenous oscillatory activity ignores the well-established fact that many oscillations are distinctly non-sinusoidal (for review, see: Cole and Voytek 2017). Acceptance of this approximation has been driven by the belief that idiosyncrasies in oscillation shape do not provide any information about underlying generative processes. However, a growing body of evidence provides evidence to the contrary. For example: the shape of delta oscillations recorded in rat motor cortex are altered following lesion of dopaminergic cells in basal ganglia (Parr-Brownlie *et al*. 2022); non-sinusoidal features of theta cycles recorded from rat hippocampus reflect synchronisation, activation sequence and firing rate of pyramidal neurons and interneurons (Cole and Voytek 2018) and have been associated with running speed (Ghosh *et al*. 2020, Quinn *et al*. 2021b); the shape of sensorimotor beta oscillations reflects thalamic input to cortical areas (Sherman *et al*. 2016, Bonaiuto *et al*. 2021), and shows increased sharpness in Parkinson’s patients, with changes correlating to limb rigidity (O’Keeffe *et al*. 2020) and being normalised by treatment (Cole *et al*. 2017, Jackson *et al*. 2019). Consequently, non-sinusoidal features of neural oscillations can index physiological information that is functionally and clinically relevant.

Although the developing literature demonstrates the utility of oscillation shape as an index of neural function, our understanding of the associated physiological processes is largely driven by invasive techniques, animal models or computational methods. In contrast, interpretation of shape within other contexts, such as human electro- (EEG) or magnetoencephalography (MEG) recordings, remains unclear. Given that these represent more commonly applied electrophysiological measures of human brain activity, this represents a significant limitation to the broader application of shape metrics. One technique that has been effective for addressing this issue is to use pharmacological intervention to modulate neurotransmission through specific neuronal receptors, and record associated changes in EEG/MEG recordings. Within this pharmaco-M/EEG approach, drug-related changes in the outcome measure suggest generative contributions from the targeted receptor. This approach has underpinned identification of the role played by key neurotransmitters (e.g., γ-aminobutyric acid [GABA] and glutamate) and ions (e.g., calcium, sodium and potassium) in several electrophysiological measures (e.g., Mucci *et al*. 2006, Muthukumaraswamy 2014, Purdon *et al*. 2015, Darmani and Ziemann 2019).

The main aim of the current study was, therefore, to use a pharmacological intervention to investigate the neurophysiological processes that contribute to the non-sinusoidal shape of neural oscillations. To achieve this, we focussed on oscillations in the alpha band (7–14 Hz), given their predominance across individuals, large signal-to-noise ratio, and well established non-sinusoidal shape. Subsequently, alpha shape was examined in healthy participants before and after oral intake of diazepam, a positive allosteric modulator of GABA type A receptors (GABA_A_) that has well established effects on alpha oscillations (Lozano-Soldevilla 2018) using a randomized placebo-controlled double-blind crossover design. Given recent evidence showing that several spatially distinct alpha oscillations can be identified based on their shape (Giehl and Siegel 2024, Bender *et al*. 2025), effects of diazepam were contrasted between alpha rhythms originating from different cortical areas.

## Methods

### Dataset and Participants

The current study utilised resting-state EEG data that were collected but not examined during a previously published project (Gordon *et al*. 2023). While experimental information relevant to the current study are reported here, the reader is referred to the previous publication for additional details about the original protocol. Recordings were collected from 21 healthy, right-handed participants aged between 18 and 50 years (mean age ± SD = 25.5 ± 4.7, 14 female). Exclusion criteria included use of centrally acting medication, history of psychiatric or neurological disease, history of alcohol or illicit drug use, and current pregnancy or breastfeeding. Written, informed consent was required prior to inclusion, all experimentation was conducted in accordance with the Declaration of Helsinki, and the study was approved by the ethics committee of the medical faculty of the University of Tübingen (approval number 456/2019BO2). As data from 6 participants failed to meet source matching and alpha SNR exclusion criteria (detailed below), a total of 15 participants were included in the analysis: 12 of these contributed data to both sessions, 3 contributed data to the placebo session only, and 3 contributed data to the diazepam session only.

### Experimental Arrangement

Participants attended two experimental sessions held at least 1 week apart, during which they sat comfortably on a reclining chair in a quiet room. Each session involved recording of eyes-open, resting-state EEG before and after oral intake of either 20 mg diazepam (diazepam; Ratiopharm) or placebo (placebo; P-Tabletten Lichtenstein). The order of diazepam and placebo sessions was pseudorandomised between participants, and both participants and experimenters were blinded to the nature of the intervention for each session. This was facilitated by the drugs having comparable appearance and being labelled with identifying codes that were only revealed following study completion. Drug intake occurred immediately following baseline EEG recording, with post-intervention EEG being collected 60 mins after drug ingestion to allow diazepam to reach peak plasma concentrations (Shader *et al*. 1984). Scalp EEG was recorded using an EasyCap EEG cap (Brain Products, GmbH, Germany), with 64 Ag-AgCl sintered ring electrodes (standard 10-10 locations) connected to a NeurOne amplifier (Bittium, Finland). Recordings were referenced to FCz, the ground electrode was positioned at PPO1h (centred between P1-Pz and PO3-POz) and impedance was maintained < 5 kΩ. All recordings were filtered on-line (0.16 Hz – 1.25 kHz) and digitised at 5 kHz prior to storage for off-line analysis. Five minutes of resting-state data were collected at each time point, during which participants fixated on a black cross displayed 1 m in front of them.

### Data analysis

#### EEG preprocessing

Data were down-sampled to 1000 Hz and line noise removed using the zap-line plugin for EEGLAB (v2024.0)(Delorme and Makeig 2004) on the Matlab platform (R2021b, MathWorks, USA). Data were then exported to MNE-Python (v1.8.0)(Gramfort *et al*. 2013, Larson *et al*. 2024), where band-pass filtering (1-100 Hz) was applied using the *filter* method for raw objects, followed by identification of bad channels using the *find_all_bads* method of the PyPrep toolbox (v0.4.3) (Appelhoff *et al*. 2025). Data segments contaminated by muscle activity or other noise were then manually identified and removed. Independent component analysis (ICA) using the FastICA algorithm (Hyvärinen and Oja 2000) was then applied to data concatenated over Pre and Post time points, separately for each session. Examination of topographies and time series was used to identify and subsequently remove components associated with blinks, saccades or ECG.

#### Spatial filtering & component identification

The conflation of source activity driven by volume conduction is an important consideration for the current study, as this spatial mixing of activity can confound interpretation of the shape of an oscillation examined in sensor space (Schaworonkow and Nikulin 2019). An approach to address this issue is to apply spatial filters that separate activity derived from different sources. Within the current study, this was achieved using an approach recently established by Bender *et al*. (2025).

This involved spectral analysis and parameterisation, application of spatio-spectral decomposition (SSD) to separate potential sources of interest (Nikulin *et al*. 2011), and source reconstruction and template matching to identify likely anatomical origins for each source. These steps are detailed below.

##### Spectral analysis and parameterisation

Pre-processed, broadband data recorded at centroparietal electrodes (C1, Cz, C2, CP1, CPz, CP2, Pz) were split into 10 s epochs prior to spectral analysis with the multi-taper method (2-40 Hz). These data were then separated into periodic and aperiodic components using the *specparam* toolbox (v1.1.0)(Donoghue *et al*. 2020). Spectral activity was modelled with peak width limits of 1 and 12 Hz, a maximum of 6 peaks, minimum peak height of 2 dB and fixed aperiodic mode. Model fit for all spectra was assessed using *r*^2^ and error values, which revealed consistently high performance (mean *r*^2^ = 98.3 ± 0.03, mean error = 0.03 ± 0.01). Individual alpha peak frequency was defined as the centre frequency for the largest peak in the 7–14 Hz range, collapsed across the 6 examined electrodes. Parameterisation was performed separately for data from each time point and session.

##### Spatio-Spectral Decomposition (SSD)

SSD decomposes the data into components with spectral power maximised within a band of interest, while minimising spectral power in flanking bands. For the current study, SSD was applied to data concatenated over Pre and Post time points, separately for each session. The band of interest was defined for each participant as the individual alpha peak frequency ± 2 Hz, averaged over Pre and Post time points within each session. The flanking frequencies were then defined as the 2 Hz bands either side of the band of interest (Nikulin *et al*. 2011). The filters estimated by SSD were then applied to the broadband-filtered data to obtain time-series for each component. Additionally, spatial patterns for each component were calculated according to Haufe *et al*. (2014). To ensure the presence of an oscillatory peak, and to identify possible slight deviations in the associated peak frequency, each time series generated by SSD was submitted to spectral parameterisation (as detailed in the previous section). Only components having an oscillatory peak in the alpha range and signal-to-noise ratio (SNR) exceeding the group-level median (3.87 dB) were considered further to ensure reliable presence of an alpha-rhythm and therefore reliable measurement of alpha waveform shape. In the final dataset, each participant contributed an average of 2.7 ± 1.4 and 2.3 ± 1.4 alpha components to the placebo and diazepam sessions, respectively (unpaired *t*-test, *P* = 0.4).

##### Source reconstruction & template matching

A forward solution was computed with the MNE *make_forward_solution* function, using a boundary element model and source space derived from the *fsaverage* template data (available as precompiled files within the MNE package), *mindist* = 5 mm, and fixed dipole orientation. The absolute cosine distance between the spatial patterns generated by SSD, and the spatial patterns from the forward model was then calculated. The component location was defined as the position where this distance was minimised, and only components fit with a distance < 0.15 were considered. These locations were then categorised according to their cortical lobe using the HCP-MMP1 parcellation (Glasser *et al*. 2016). As only a single source was localised to frontal cortex, data from this area was not considered in the analysis.

#### Waveform shape analysis

The shape of oscillatory activity within the time series extracted from each SSD component was quantified using a previously established pipeline, involving application of empirical mode decomposition (EMD) to derive narrow band modes in the alpha range, followed by calculation of phase-aligned instantaneous frequency to describe shape of individual oscillatory cycles (Quinn *et al*. 2021b, Opie *et al*. 2024). To reduce the impact of mode mixing (Huang *et al*. 1999, Deering and Kaiser 2005), the masked version of EMD was applied using the *mask_sift* function of the EMD toolbox (v0.8.1) (Quinn *et al*. 2021a). A maximum of 6 intrinsic mode functions (IMFs) were generated using masking frequencies of 120, 64, 32, 11, 7 and 2 Hz. All subsequent analyses focussed on the alpha mode. First, fit quality was assessed via visual inspection compared with the raw SSD component time series. Individual alpha cycles were then identified based on the instantaneous amplitude and phase of the signal, derived using the normalised Hilbert transform. To avoid potentially confounding effects of variations in amplitude, only large amplitude cycles were included (top 25^th^ percentile). Cycles were additionally required to have unique control points (i.e., ascending/descending zero crossing, peak, trough; Quinn *et al*. 2021b) and not show phase reversals. Using these criteria, a total of 61,311 cycles were identified. Instantaneous frequency (IF) of identified cycles was then quantified as the first derivative of the instantaneous phase with respect to time (Huang *et al*. 2009). To facilitate comparison of cycles across consistent features (i.e., ensure peaks are compared with peaks), IF values were then projected to a common phase space, producing phase aligned IF (IF*_PA_*) values. To characterise variation in alpha shape across individual cycles, IF*_PA_* values were subsequently decomposed using principal component analysis (PCA). The resulting principal components (PCs) provide a series of waveform motifs that each describe major sources of variance relative to the mean IF profile, with larger scores indicating shapes that are further from the mean. PCA was applied using the *sails* toolbox (v 1.7.0)(Quinn and Hymers 2020) on demeaned data that was concatenated over subjects and sessions. The first four PCs explained >95% of variance in the data and were therefore used for subsequent statistical analysis.

### Statistical analysis

#### Spectral data

Parameterised spectral data (aperiodic slope, alpha peak frequency and power) averaged over centroparietal electrodes was compared between sessions (diazepam, placebo) and time points (Pre, Post) using Bayesian generalised linear mixed models (GLMM). For alpha power and peak frequency data in the diazepam session, one participant was missing data in the Post time point, whereas another participant was missing data at Pre and Post time points. Consequently, these spectral data were imputed prior to statistical analysis with multiple imputation (20 imputed datasets) using the *mice* package (Van Buuren and Groothuis-Oudshoorn 2011). Data were modelled with a skew-normal distribution and identity link function, with maximal by-participant random effects for time (model 1). Each model was run using 8 independent chains, with 1000 warm up and 3000 post-warm up samples (totalling 480,000 post warm-up samples across the 20 imputed datasets), and default flat priors.

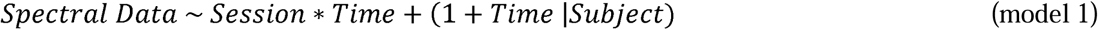

#### Shape data

Bayesian GLMMs were also used to assess changes in alpha shape over time (Pre, Post) and between cortical lobes (Occipital, Temporal, Sensorimotor, Parietal). A single model was run for each PC of interest: each model incorporated all data that met the quality control measures defined above, and used a student’s *t* distribution with identity link function. However, between-session variance in the presence of some oscillatory activity meant it was not possible to match SSD components between sessions. Consequently, the specific participant cohort included within each session was not matched, and we were unable to make direct comparisons between data from the placebo and diazepam sessions. In an attempt to address this limitation, the model implemented maximal by-participant random effects for both time and SSD component (model 2). Nonetheless, within-session contrasts were sufficient to address the study’s main aims. Each model was run using 4 independent chains, with 1000 warm up and 3000 post-warm up samples (totalling 12,000 post warm-up samples), and weakly informative priors (Normal (0, 1)) for fixed effects.

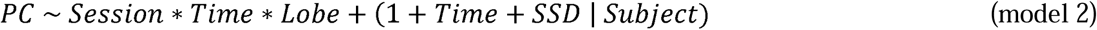

To further investigate drug-related shifts in the distribution of PC scores, additional comparisons across Pre and Post time points were made using two-sample Cramér-von Mises tests, implemented in the *SciPy* toolbox (v1.14.1)(Virtanen *et al*. 2020). A Bonferroni correction was applied to adjust for multiple comparisons in these data, with P < 0.003125 (i.e., 0.05 / [4 principal components * 4 cortical lobes]) considered significant.

#### Bayesian estimation

All GLMMs were computed in R (v 4.4.3) using RStudio (v2024.12.1). Posterior distributions were estimated within BRMS (Bürkner 2017), using the No-U-Turn Sampler (NUTS) extension of Hamiltonian MCMC. Chain convergence was assessed using Rhat values < 1.1 and posterior predictive checks (Gelman and Rubin 1992, Gabry *et al*. 2019). Custom contrasts of main effects and interactions were generated using the *emmeans* package (Lenth 2025), with effect existence and significance identified using the probability-of-direction (e.g., pd; Makowski *et al*. 2019) and region of practical equivalence (ROPE), respectively. For the current study, ROPE was defined as ± 5% of the standard deviation for the data being compared (Kruschke 2018). The null hypothesis of no difference was accepted for comparisons where the 89% highest density interval (HDI) fell completely inside the ROPE (i.e., 100% in ROPE). In contrast, the null hypothesis was rejected when the 89% HDI fell completely outside the ROPE (i.e., 0% in ROPE). No decision was made if the 89% HDI partially overlapped ROPE (Kruschke 2018, Opie *et al*. 2024, Moore *et al*. 2025). Results for all Bayesian models are presented as median values [lower 89% HDI, upper 89% HDI].

## Results

### Drug-related changes in spectral data

First, to align our results to previous literature, we compared spectral characteristics between sessions and time points (Figure 1). At baseline, between-group comparisons for all measures were inconsistent and failed to provide sufficient evidence to accept or reject the null hypothesis (all *pd* < 80%, % in ROPE: 29.7 – 41.7%). For all measures in the placebo session, in addition to alpha frequency in the diazepam session, within-group comparisons were also inconsistent and failed to provide sufficient evidence to accept or reject the null hypothesis (all *pd* < 86%, % in ROPE: 21.0 - 44.9%). In contrast, diazepam was associated with consistent and significant reductions in both aperiodic slope (*pd* = 99%, 0% in ROPE; Fig 1B, right) and alpha power (*pd* = 99%, 0% in ROPE; Fig 1C, right) at the Post time point.

**Figure 1.**
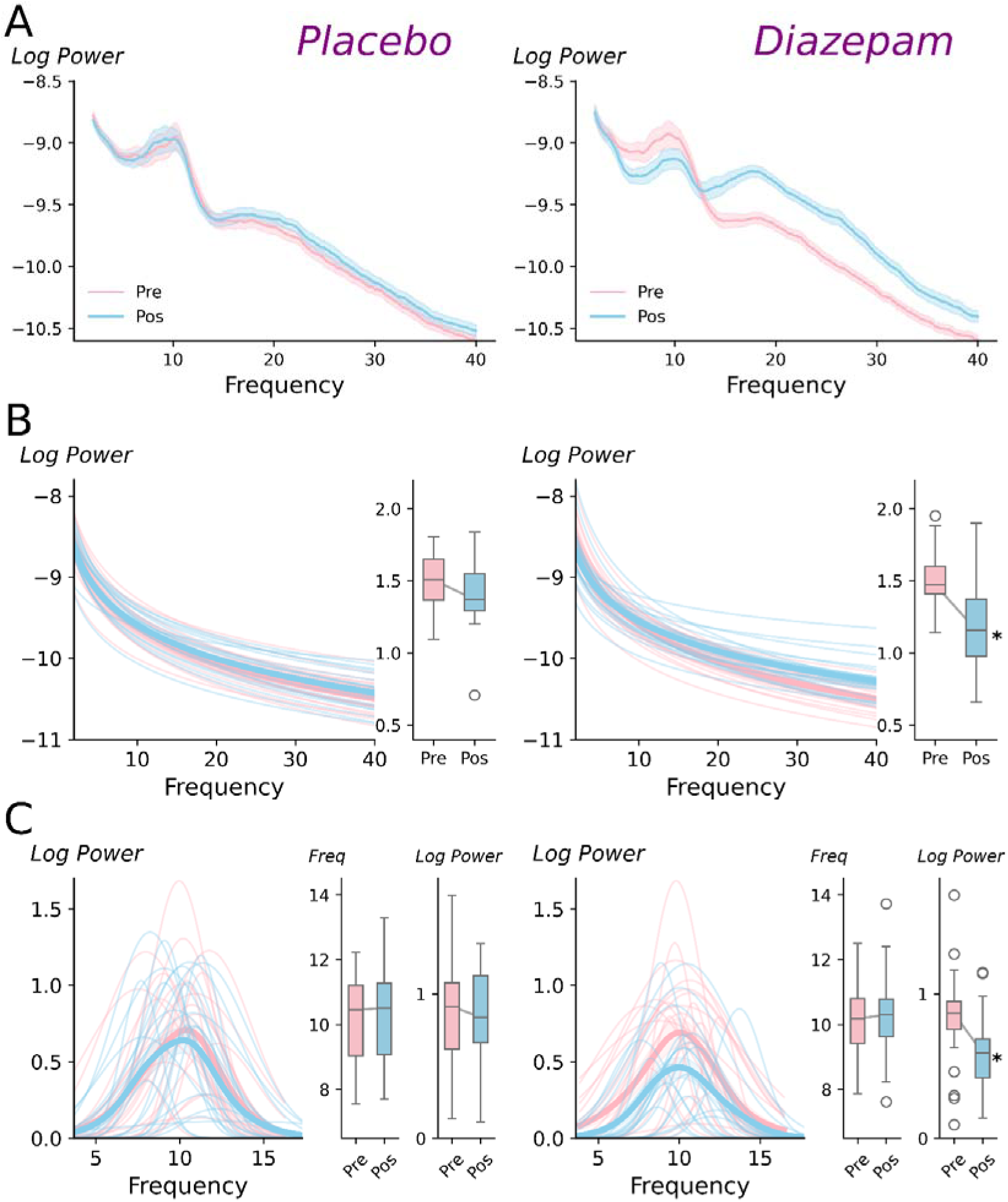
Effects of diazepam and placebo on periodic and aperiodic spectral features. ***(A)*** Power spectral density curves (log(V^2^/Hz) recorded before *(Pre)* and after *(Post)* intake of placebo *(left)* or diazepam *(right)*. Shaded area shows +/- 1 standard error of the mean ***(B)*** Aperiodic components *(line plot)* and slopes *(box plot)* plotted before *(Pre/ pink)* and after *(Post/ blue)* intake of placebo *(left)* or diazepam *(right)*. ***(C)*** Periodic components *(line plot)*, alpha peak frequency *(middle box plot)* and alpha power *(right box plot)* recorded before and after placebo (*left*) or diazepam (*right*). For panels B & C, thick lines show group average and thin lines show responses for individual participants. Furthermore, lower, middle and upper lines of boxplots indicate 25^th^, 50^th^ and 75^th^ percentiles, respectively, whiskers extend to 1.5 x the interquartile range. \**pd* > 99%, 0% in ROPE, relative to *Pre*.

### Drug-related changes in alpha waveform shape

Having established directionality of changes in spectral measures, we now turn to waveform shape analyses: for each cortical area of interest, Figure 2 shows IF*_PA_* profiles recorded before and after placebo and diazepam, in addition to the spatial patterns of the underlying alpha components.

**Figure 2.**
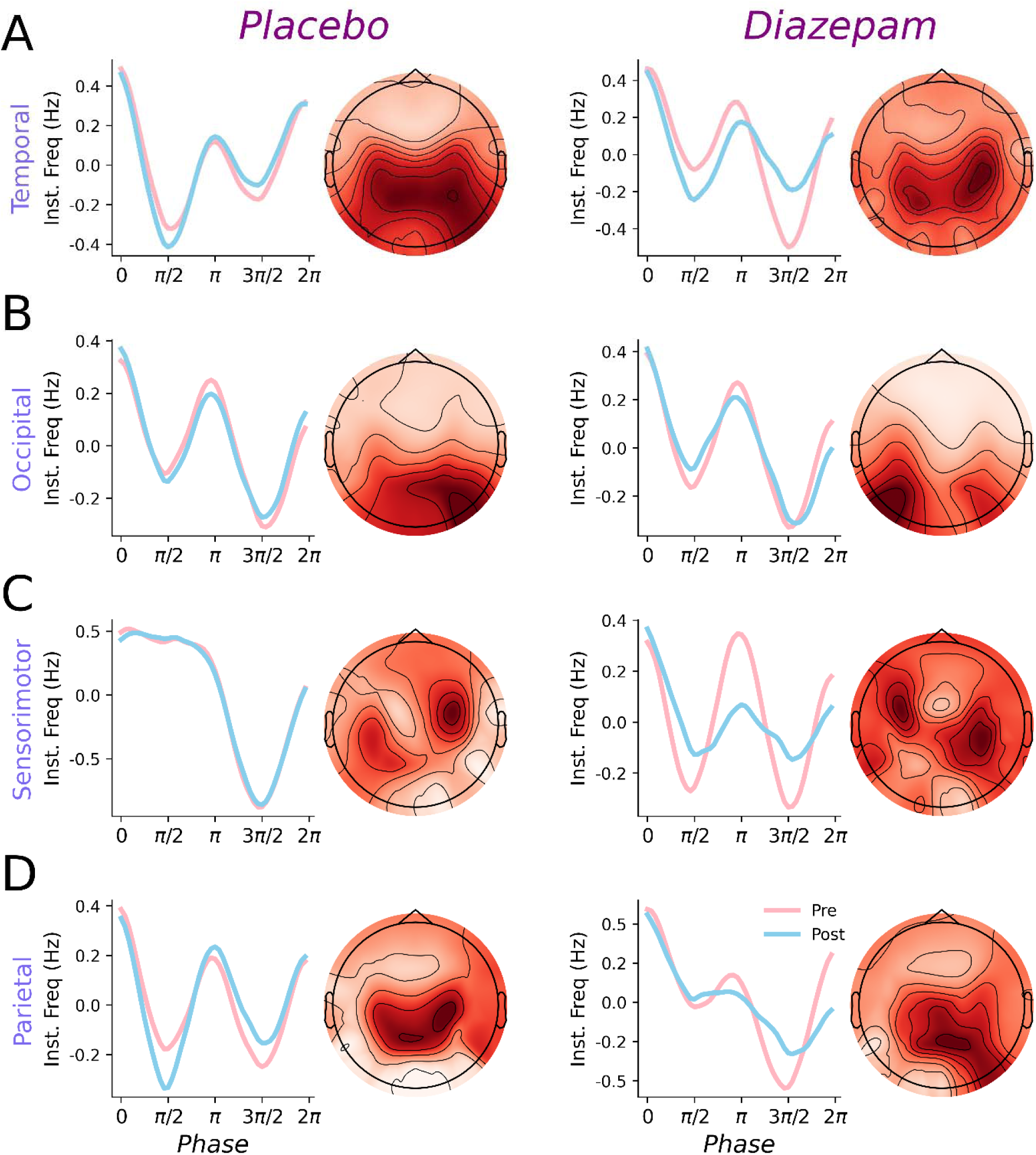
Effects of diazepam and placebo on IF*_PA_*profiles in each cortical area. Group-averaged IF*_PA_* profiles *(line plots; pre in pink, post in blue)* and spatial patterns associated with the included SSD components *(topoplots)* for temporal ***(A)***, occipital ***(B)***, sensorimotor ***(C)*** and parietal ***(D)*** cortices. Responses from the placebo session are shown in the left column, whereas responses from the diazepam session are shown in the right column.

Across areas, IF*_PA_* profiles tended to indicate that alpha peaks and troughs were associated with reduced frequency, whereas zero crossings were associated with increased frequency. To quantify these changes, PCA was used to decompose these profiles into waveform motifs, with the first 4 components being characterised based on their IF*_PA_* and normalised waveforms in Figure 3. These components explained more than 95% of variance in the data, although the majority was partitioned within components characterising peak/trough (PC1, ∼43%) and edge (PC2, ∼38%) asymmetry.

**Figure 3.**
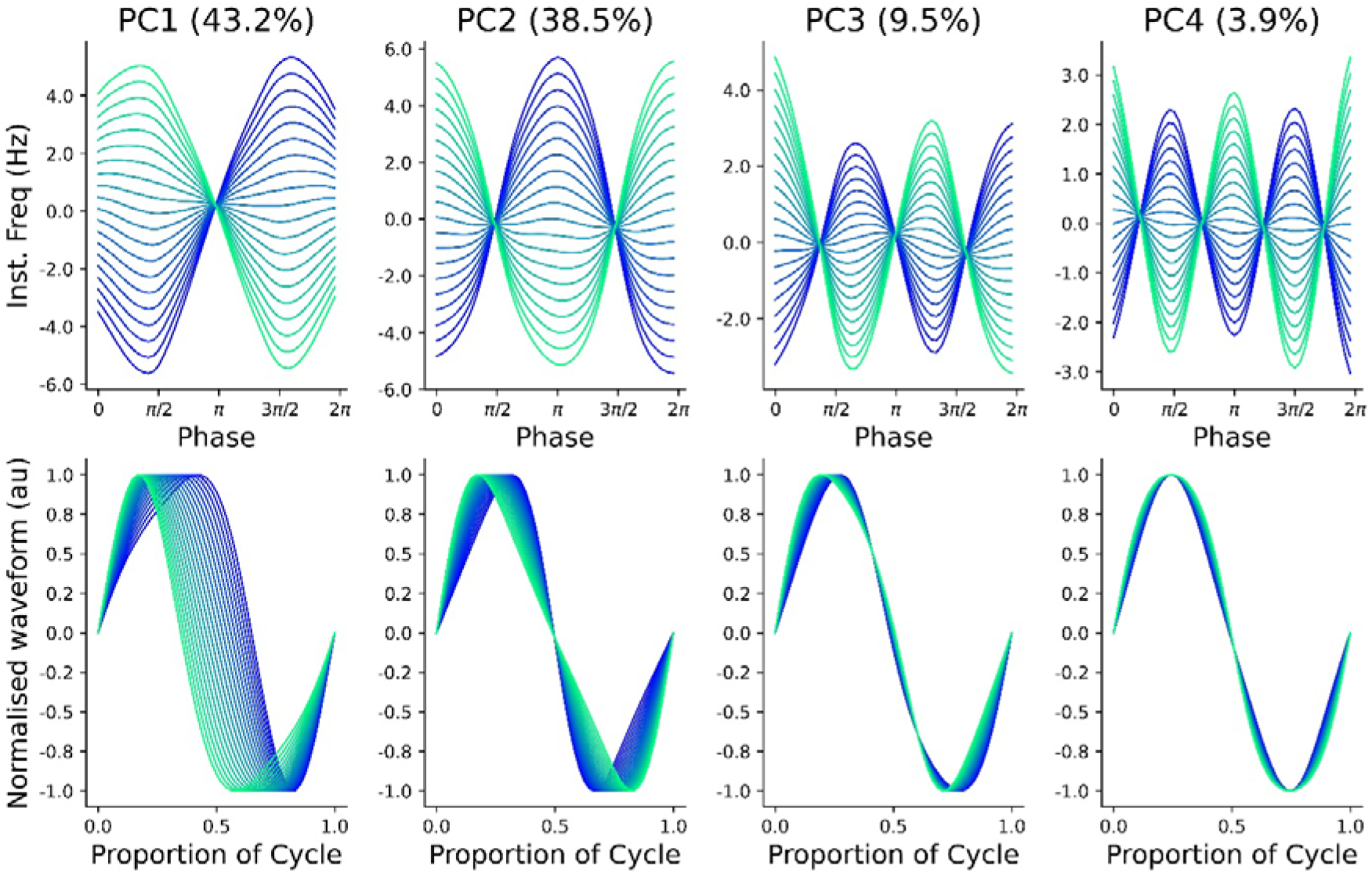
Waveform motifs identified with PCA. IF*_PA_* profiles *(top row)* and associated normalised waveforms *(bottom row)* for the top four principal components (PC, percentage indicates proportion of variance explained), plotted across the percentiles of scores observed for all alpha cycles. Colour gradient ranges from 1^st^ *(blue)* to 99^th^ *(green)* percentile.

To quantify drug-related changes in alpha shape, scores for the first 4 PCs were compared between time-points, with results plotted separately for each cortical area in Figure 4A-4C and supplementary Figure S1. Diazepam was associated with consistent and significant reductions in scores for PC1 in temporal cortex (*pd* = 99%, 0% in ROPE; Fig 4A), and PC4 in temporal (*pd* = 99%, 0% in ROPE), sensorimotor (*pd* = 100%, 0% in ROPE) and parietal (*pd* = 100%, 0% in ROPE) cortices (Fig 4C). Furthermore, consistent and significant increases in PC3 scores were found in both temporal (*pd* = 100%, 0% in ROPE) and parietal cortices (*pd* < 99%, 0% in ROPE; Fig 4B). While comparisons of PC4 in occipital cortex also showed consistent changes after diazepam (pd = 97.8%) this failed to reach a practical level of significance (8.8% in ROPE). All other comparisons were inconsistent and failed to provide sufficient evidence to accept or reject the null hypothesis (all *pd* < 96.6%, % in ROPE: 16.6 – 87.8%; Fig S1). To illustrate how diazepam-related changes in PC scores influenced alpha shape, normalised waveforms associated with the median score for each component are plotted in Figure 4D.

**Figure 4.**
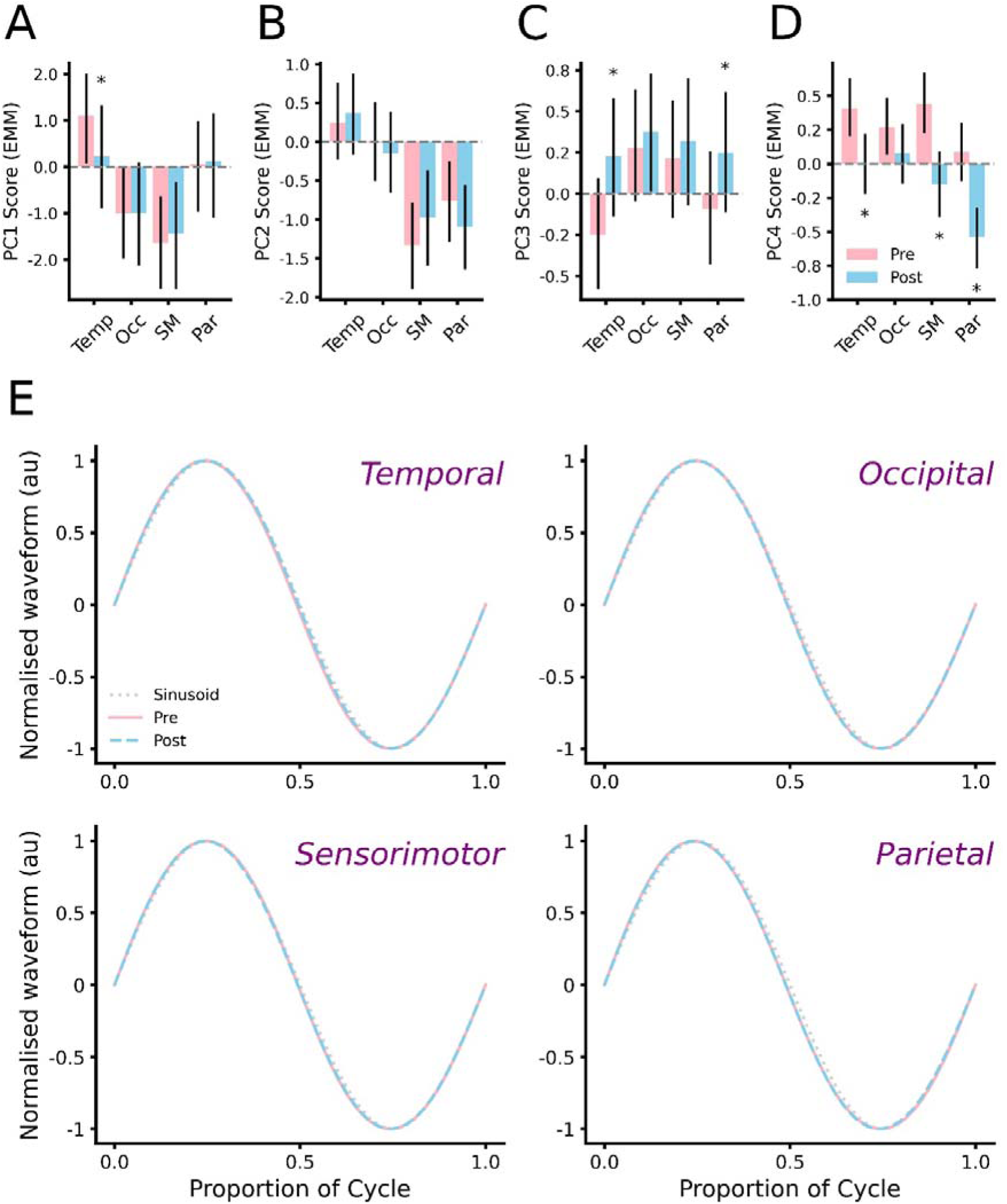
PC scores identify subtle changes in alpha shape. ***(A-D)*** Estimated marginal means from Bayesian GLMMs showing mean scores for PC1 *(A)*, PC2 *(B),* PC3 *(C)* and PC4 *(D)* before *(pink)* and after *(blue)* diazepam. Temp: temporal, Occ: occipital, SM: sensorimotor, Par: parietal. Error bars show 89% HDI. ***(D)*** Normalised waveforms associated with median PC scores for each cortical area recorded before *(pink curve)* and after *(blue curve)* diazepam. Sinusoid plotted for reference *(grey dotted curve)*. \**pd* > 99%, 0% in ROPE, relative to *Pre*.

We examined the changes in score distribution across Pre and Post conditions via two-sample distributional testing for each PC. To facilitate data interpretation, distributions showing significant drug effects are plotted in Figure 5, whereas all other distributions are plotted in supplementary figures S2 and S3. For the diazepam session, there were significant differences in score distribution at the post time point for PC1 (ω^2^ = 0.96, *P* = 0.003), PC2 (ω^2^ = 1.41, *P* = 0.0003) and PC3 (ω^2^ = 1.35, *P* = 0.0004) in sensorimotor cortex (Fig 5A), in addition to PC1 in parietal cortex (ω^2^ = 1.38, *P* = 0.0004)(Fig 5B). For these components, changes in distribution consistently involved an increase in more extreme scores (i.e., bottom and top 10^th^ percentile). Normalised waveforms associated with these more extreme scores are shown in Figure 5C and 5D. All other comparisons failed to show any significant change in distribution (Fig S2 & S3).

**Figure 5.**
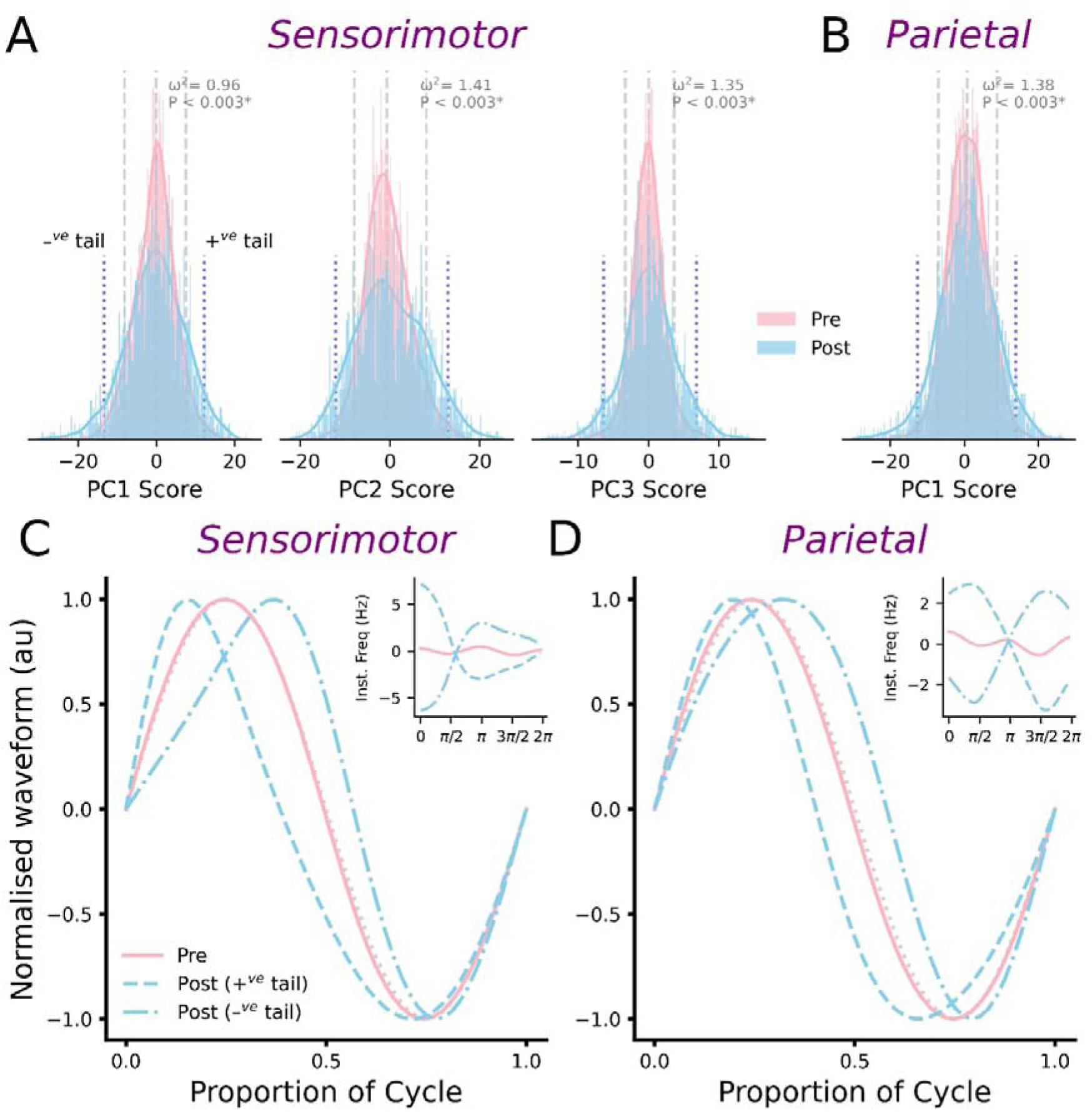
Extreme alpha shapes are increased after diazepam. ***(A-B)*** Probability densities for PC scores from sensorimotor *(A)* and parietal *(B)* cortices, which showed significant changes in distribution due to diazepam. Vertical dashed grey lines indicate 10^th^, 50^th^ and 90^th^ data percentiles. Post-diazepam alpha tended to have an increase in scores in the top and bottom 10^th^ percentiles. ***(C-D)*** Normalised waveforms associated with diazepam-related changes in score distribution. Waveforms were derived using scores identified by vertical dotted blue lines in panels A and B (denoted *+^ve^ tail* and *–^ve^ tail* in panel A). For PCs failing to show significant change in distribution, scores were set at the median value. Inset shows IF*_PA_* profiles for each curve.

## Discussion

While examination of neural oscillations generally focusses on spectral features such as power, frequency and phase, a developing literature has identified oscillation shape as providing additional neurophysiological insight. However, the link between oscillation shape and specific neural mechanisms such as neurotransmission has not been established in non-invasive human recordings. Within the current study, we began to address this underexplored field by investigating changes in the shape of alpha oscillations induced by potentiation of GABA_A_-mediated inhibitory neurotransmission. This was achieved by quantifying alpha shape derived from resting-state EEG recordings before and after intake of diazepam or placebo. Using this approach, an initial analysis found significant changes in PCs that described subtle shifts in alpha shape, the specific nature of which varied between cortical areas. However, additional analysis focussing on alterations in the distribution of PC scores identified stronger effects of diazepam that were limited to alpha cycles originating from sensorimotor and parietal cortices. These results provide the first evidence that, when assessed non-invasively with EEG, GABA_A_ergic processes can be indexed via the shape of alpha oscillations.

### Spectral effects of diazepam

Prior to waveform analysis, effects of increasing neurotransmission through GABA_A_ receptors were examined using a conventional spectral decomposition technique that involved Fourier methods (i.e., the multi-taper method). Consistent with findings from previous studies (for review, see; Lozano-Soldevilla 2018), this analysis identified a significant reduction in alpha power without any change in frequency (Fig 1). These effects have been suggested to reflect drug-related changes in neuronal firing that do not influence oscillatory frequency (e.g., reduced spikes per burst without a change in burst frequency), or alterations to thalamic input to cortical alpha generators (Lozano-Soldevilla *et al*. 2014). Irrespective of the specific mechanism, reductions in alpha power demonstrate that diazepam was effective at producing the expected changes in oscillatory activity. However, we additionally observed a drug-related reduction in the aperiodic component, corresponding to a flattening of the PSD (Fig 1). Previous simulation work has suggested that the aperiodic slope is an index of the balance between neural excitation and inhibition (Gao *et al*. 2017); while the drug-related changes observed here are consistent with this idea, the direction of change was not. Specifically, current interpretations of aperiodic slope suggest potentiation of inhibitory function should drive an increase in slope, as opposed to the reduction we found. However, it is important to note that the nature of the relationship between aperiodic activity and neural activity is still being clarified (Donoghue 2024). In particular, recent studies using pharmacological intervention or ethanol consumption to potentiate GABAergic function have reported both increased (Stock *et al*. 2020, Waschke *et al*. 2021, Gonzalez-Burgos *et al*. 2023, Salvatore *et al*. 2024, Barone *et al*. 2025) and decreased (Barone *et al*. 2025) aperiodic activity. This includes contradictory responses to the same drug (i.e. diazepam; current study *vs.* Gonzalez-Burgos and colleagues) and drug class (i.e., positive allosteric modulators of GABA_A_ receptors; current study *vs.* Barone and colleagues). While methodological differences possibly contributed to this variability, these outcomes nonetheless demonstrate that further work is needed to clarify the neurophysiological interpretation of aperiodic activity.

### Diazepam alters the shape of alpha oscillations

To examine drug-related changes in alpha shape, the current study applied a recently developed waveform analysis pipeline (Quinn *et al*. 2021b). This involved derivation of IF*_PA_* profiles for individual cycles, and subsequent decomposition with PCA to extract motifs describing key features (e.g., Figs 2 & 3). Consequently, changes in PC scores over time can be interpreted as changes to specific facets of alpha shape. Using this approach, we found that scores for the top four PCs were not altered in the placebo session, and this was consistent across all cortical areas. In contrast, diazepam was associated with significant changes in PC1, PC3 and PC4, the nature of which differed between cortical areas: while changes in PC1 were limited to temporal cortex, effects on PC3 were apparent in both temporal and parietal cortices, whereas PC4 was altered in all cortices other than occipital (Fig 4A-4C). Taken together, these results demonstrate that examination of oscillation shape can reveal regionally specific effects of GABA_A_-mediated inhibitory neurotransmission on alpha cycles. However, while these changes were significant, it is important to note that their impact on shape was highly subtle. This is demonstrated in the normalised waveforms plotted for each cortical area in Figure 4D. Within these, all waveforms show subtle deviations in shape with respect to a sinusoid which are consistent with the known non-sinusoidal shape of alpha cycles (Cole and Voytek 2017). While differences between Pre and Post waveforms are only modest, it is important to note the macroscale nature of the EEG recordings from which these data were derived. Consequently, these modest differences do not necessarily preclude changes in neurophysiological dynamics.

Despite this, the ‘shape space’ described by each PC is large and variable. Furthermore, recent work demonstrates that functionally significant changes in shape may be limited to sub-populations of cycles. Specifically, performance of reaching movements was associated with changes in the shape of beta oscillations that were limited to cycles having more extreme shape (i.e., scores furthest from the median)(Rayson *et al*. 2023, Szul *et al*. 2023). Consequently, the possibility remained that different effects of diazepam may be apparent within specific sections of the score distribution. In support of this, subsequent inspection of score distributions identified several significant changes following diazepam that were not apparent with placebo. In contrast to our primary analysis, these were specific to cycles originating from sensorimotor and parietal cortices, involved a different collection of PCs, and an increase in the proportion of cycles within the tails of the distribution (i.e., more extreme shape; Fig 5A & 5B). Taken together, these results suggest that, although waveform shape can index important neuromechanistic processes, the way in which this is quantified needs further consideration. In particular, much greater exploration of different segments of the ‘shape space’ is required. An important consideration in achieving this will be to consider alternative decomposition approaches that might allow more granular description of the hugely variable ‘shape space’. Elegant examples of how this might be achieved were recently reported by work using non-linear dimensionality reduction (Lee *et al*. 2021, Sebastian *et al*. 2023) in conjunction with clustering techniques (Lee *et al*. 2021). Investigating if similar approaches have utility in non-invasive human recordings will be an exciting area for future work.

### Spatially specific effects of diazepam on alpha shape

Although sensorimotor and parietal cortices both demonstrated an increase in alpha cycles with more extreme shape following diazepam, the specific changes in shape differed between areas: while effects on sensorimotor cycles included changes in peak width and edge speed, effects on parietal cycles were more specific to alterations in peak-trough asymmetries (Fig 5C & 5D). One reason for this differential response to diazepam could be that it reflects unique features of the neural circuits generating alpha oscillations in each area. Indeed, recent work using both animal and human recordings has suggested this as an explanation for regional variations in oscillation shape (García-Rosales *et al*. 2024, Giehl and Siegel 2024). Given the pharmacological intervention we applied, variations in GABA_A_ receptors are a potential contributor here; although they are located throughout the brain, GABA_A_ receptors are highly heterogeneous in nature. In particular, there are four GABA_A_ receptors that are sensitive to diazepam; these are defined by different α subunits (α1-3 & 5), the presence of which determines unique anatomical and subcellular distribution, in addition to electrophysiological and functional effects (Sieghart and Sperk 2002, Olsen and Sieghart 2009, Engin *et al*. 2018). For example, while receptors having an α1-3 subunit tend to be located synaptically, those with an α5 subunit are more focussed on extrasynaptic membranes (Engin *et al*. 2018). Consequently, diazepam can be expected to have some influence on both phasic (i.e., synaptically mediated) and tonic (i.e., extrasynaptically mediated) inhibition (Engin *et al*. 2018), and spatial variance in this effect could contribute to the regional modulation of shape observed in the current study. Future pharmacological investigation targeting alternative GABA_A_ receptor subtypes – for example by testing zolpidem, a positive modulator mainly at the α1-GABA_A_ receptor – could be an interesting way to further investigate this concept. Modulation of other key neurotransmitter and neuromodulatory systems will also be important. In addition to characteristics of the local generative circuitry, sensorimotor and parietal areas also have unique connectivity networks (e.g., Jann *et al*. 2010), idiosyncrasies of which may contribute to regional effects of diazepam on alpha shape. In support of this, recent work reported differential changes in resting-state functional magnetic resonance imaging (fMRI) connectivity in sensorimotor and parietal cortices following administration of alprazolam (Wein *et al*. 2024), an alternative positive allosteric modulator of GABA_A_ receptors.

### Limitations

The current study includes some limitations that warrant discussion. First, while a spatial filtering technique was applied to localise different alpha sources, it is important to note that this did not use individualised structural MRI data. As we refrained from fine grained spatial interpretation of the data, potential impact of source localisation error on the reported outcomes should not be substantial. However, spatial information should nonetheless be interpreted with caution. Second, while attempts were made to match alpha sources between sessions within each subject, the nature of the spatial filtering we applied meant this was not possible for the majority of participants. While there was substantial overlap in the participants contributing data to each session (n = 12), our analysis and interpretation of the data was limited to within-subject comparisons. In an attempt to address this limitation to some extent, the GLMMs we implemented included maximal by-participant random effects (i.e., intercepts and slopes) for time and SSD component. However, future work using alternative approaches that allow sources to be matched more specifically will be important for confirming the findings we report here.

In conclusion, the current study used a pharmaco-EEG approach to investigate the extent to which the shape of alpha oscillations, recorded non-invasively in humans using EEG, can index inhibitory neurotransmission involving GABA_A_ receptors. Subsequently, while all facets of alpha shape were unchanged in the placebo session, demonstrating methodological consistency, ingestion of diazepam – a positive allosteric modulator of GABA_A_ receptors – was associated with significant changes in alpha shape. Importantly, the specific effects of diazepam on alpha shape varied between cortical areas, and included a combination of highly subtle and more clearly discernible effects. These findings provide further support for the idea that oscillation shape is neurophysiologically informative. However, they additionally highlight the need for greater investigation of the ‘shape space’, particularly with respect to identifying the functional and physiological relevance contained within different clusters of the waveform shape space.

## Supplementary Figures

**Fig S1.**
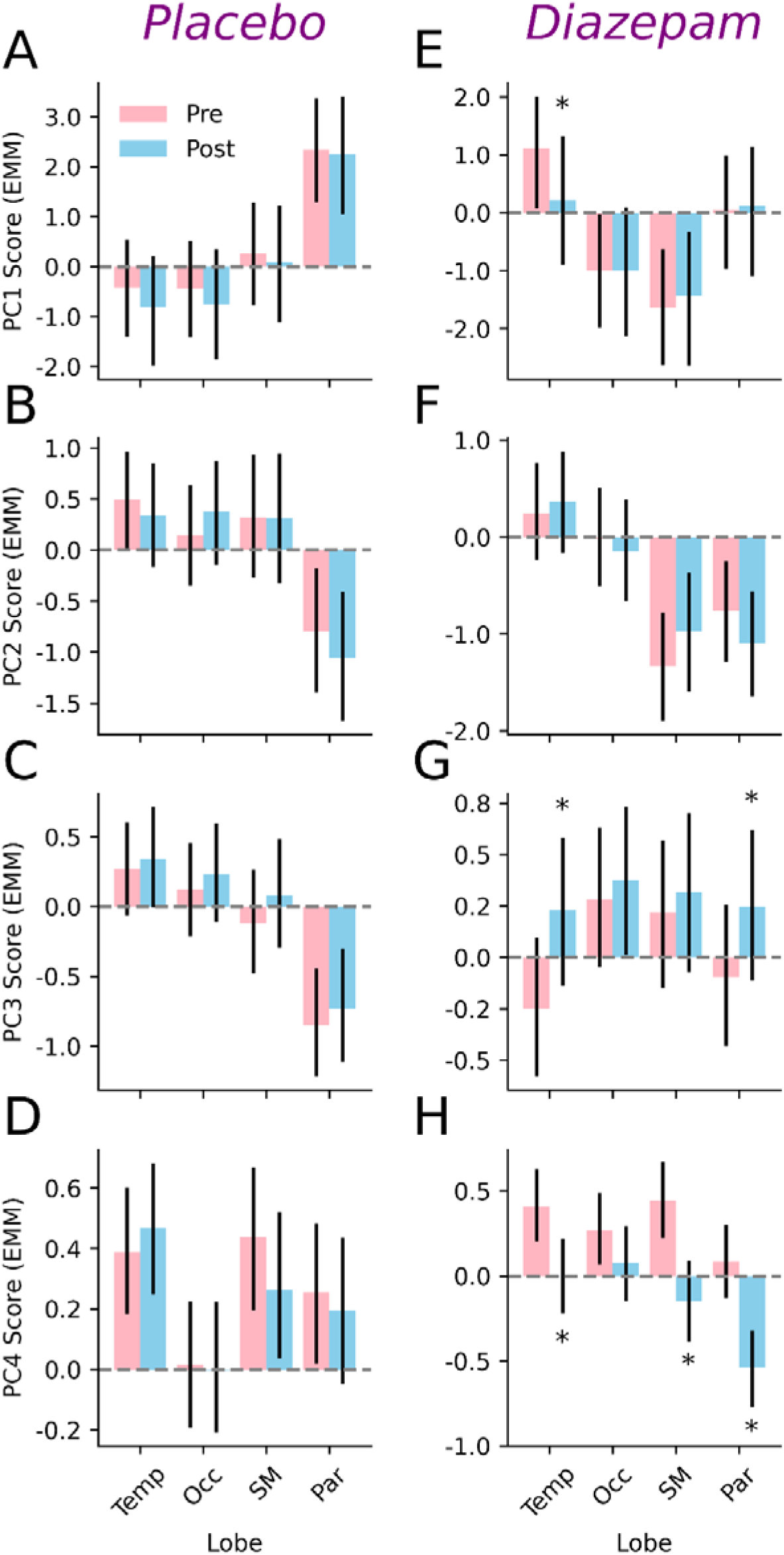
Effects of diazepam and placebo on PC scores. Estimated marginal means from Bayesian GLMMs for top four PCs *(rows)* before *(pink bars)* and after *(blue bars)* placebo ***(A-D)*** or diazepam ***(E-H)***. *pd > 99%, 0% in Rope, compared to *Pre*.

**Fig S2.**
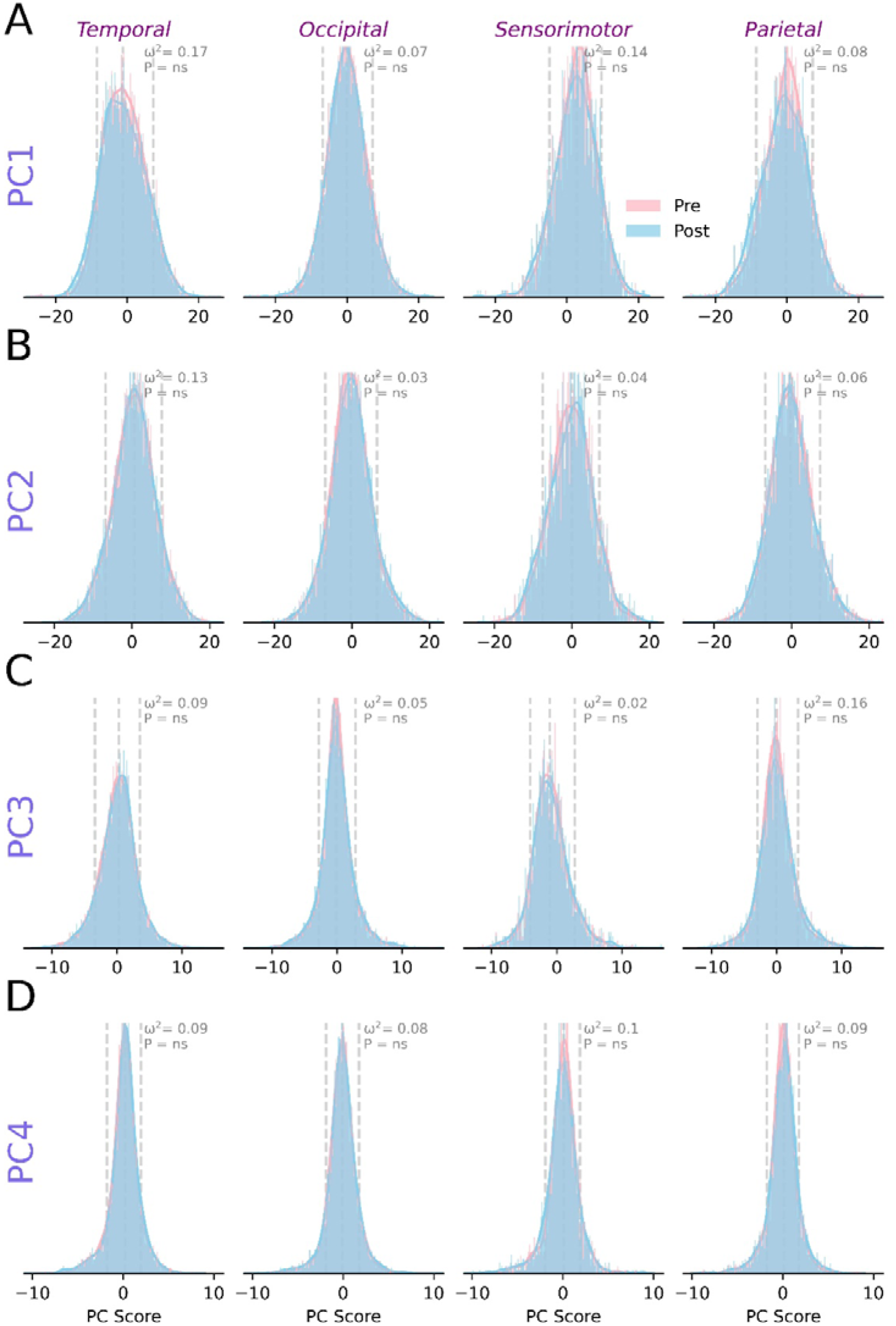
Distribution of PC scores for placebo session. ***(A-D)*** Probability densities for top four PCs before *(pink)* and after *(blue)* intake of placebo, separated by each cortical area *(columns)*. *ns =* not significant, relative to *Pre*.

**Fig S3.**
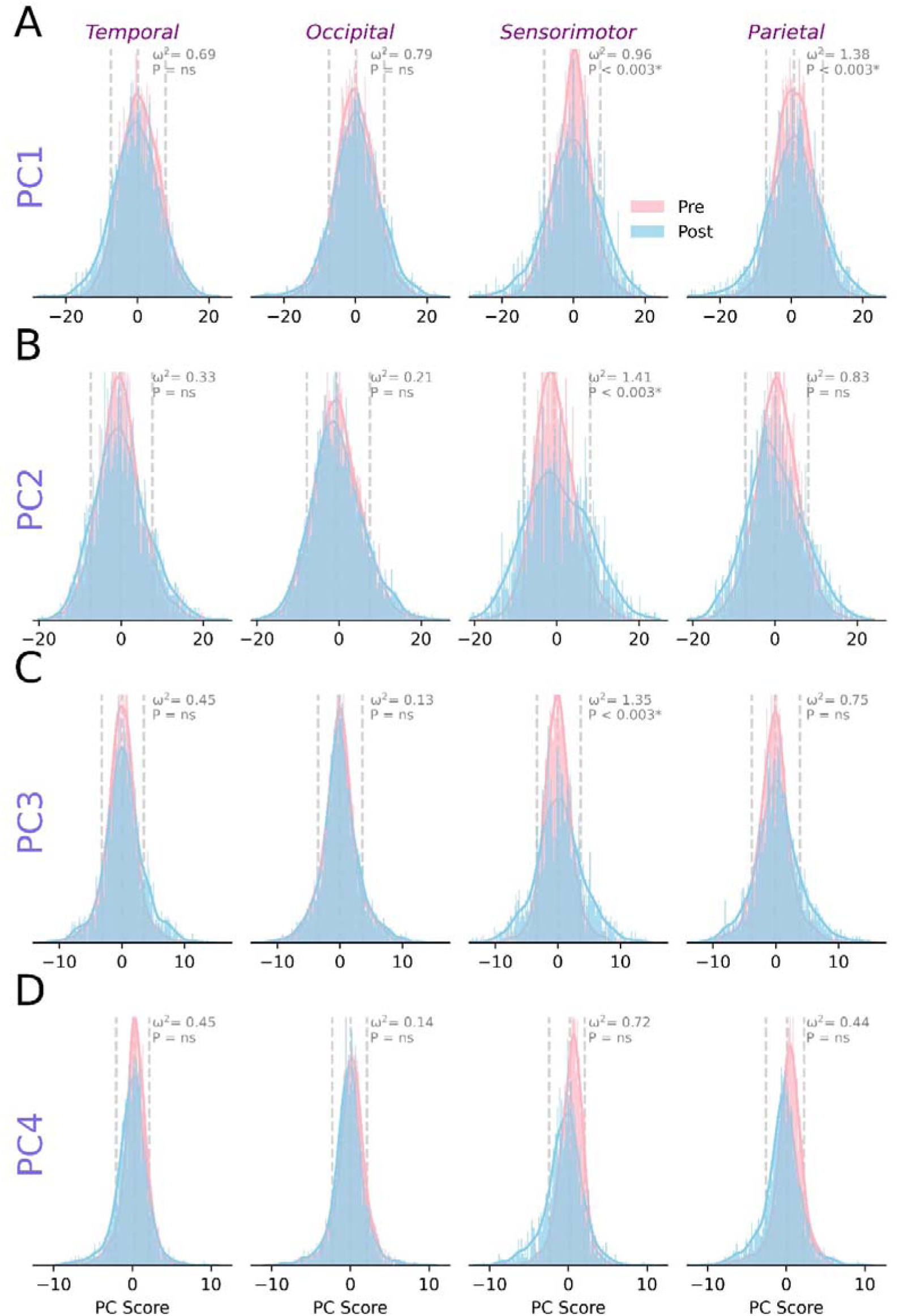
Distribution of PC scores for diazepam session. ***(A-D)*** Probability densities for top four PCs before *(pink)* and after *(blue)* intake of diazepam, separated by each cortical area *(columns)*. *ns =* not significant, relative to *Pre*.

